# Exosomal cargo genes as biomarkers and potential mediators of exosome-driven cell communication in Pleural Mesothelioma

**DOI:** 10.1101/2025.04.04.647264

**Authors:** Agnieszka Kraft, Andrea Tönz, Fabian Schläpfer, Manuel Ronner, Vanessa Orlowski, Michaela B Kirschner, Emanuela Felley-Bosco, Julia Bein, Peter J Wild, Valentina Boeva, Isabelle Opitz, Mayura Meerang

## Abstract

Pleural mesothelioma (PM) is a rare yet aggressive and heterogeneous cancer type with very poor survival rates. Due to its long latency period and nonspecific symptoms, the disease is usually detected at advanced stages, limiting available treatment options and leading to poor survival rates. So far, the disease can only be confirmed through invasive thoracoscopic biopsy. Moreover, the proposed circulating protein biomarkers lack sensitivity and specificity, highlighting the urgent need for novel approaches.

In our previous work, we characterized the transcriptomic profile of extracellular vesicles secreted by PM cells and demonstrated their feasibility as a source of circulating biomarkers. To further investigate the role of circulating RNA in tumor progression and its potential use in non-invasive testing for PM, here we present a detailed characterization of RNA cargo carried by PM exosomes - a distinct subset of extracellular vesicles known to serve as key mediators of intercellular communication. By utilizing primary cell cultures established from tissue samples of 11 PM and 6 non-PM patients, followed by exosome isolation, and total RNA sequencing of RNA from isolated exosomes, matching cells, and tissues, we provide new evidence on exosomal RNA secretion, regulation of PM-exosome cargo genes, and their potential functions in recipient cells. We show that PM-exosomal cargo is enriched in genes associated with proliferation, whose key transcriptional factors are *SETDB1*, *FOXM1* and *GATA2*. We identified Pcbp2, Srsf1 and Srsf9, RNA-binding proteins, which may be involved in selective cargo sorting into PM-exosomes. Our analysis identified six secreted genes, including *GAS5*, *AL031666.3*, *RSLD1D1*, *AC103740.2*, *ADAM10*, and *AC020892.1*, whose exosomal expression was associated with patient survival, as promising biomarkers for patient diagnosis, stratification, and prognosis assessment. Finally, we identify potential target genes of candidate LncRNAs biomarkers and processes in which they are involved.

## Background

Pleural mesothelioma (PM) is a rare yet aggressive cancer type with very poor survival rates, ranging between 9 to 17 months after diagnosis [1–4]. The disease, associated with prior asbestos exposure in most cases [5–7], develops in the mesothelial cells composing the pleura, over a span of 30 to 40 years without manifesting symptoms [8]. This often leads to disease detection only in advanced stages and limits available treatment options. Currently standard multimodal approaches that integrate surgery, chemotherapy, and radiation, remain ineffective in many cases [8–12]. Novel approaches, including ipilimumab and anti-PD-L1 antibody, nivolumab, showed promising outcomes compared to chemotherapy, but only in patients with non-epithelioid tumors [13,14]. The difficulty of treating the disease is additionally influenced by its high molecular heterogeneity. Tumors are traditionally classified into epithelioid, sarcomatoid, or biphasic subtypes, based on the dominance of cancer cells with epithelioid or sarcomatoid morphology [15]. Of note, recent studies have shown that the histological classification does not fully explain PM heterogeneity, with intratumor transcriptional heterogeneity scores, captured by the continuum of *S-* and *E-*component marker genes expression, better capturing patient clinical characteristics [16–20].

At the same time, PM lacks non-invasive and effective diagnostic tools. Currently, non-invasive disease detection is limited to imaging-based approaches, which lack sensitivity, hence a definitive diagnosis needs to be confirmed by thoracoscopic biopsy [21,22]. Despite being very sensitive, the biopsy is an invasive procedure, not applicable for routine early screening of patients. Therefore, non-invasive and robust detection methods for PM are urgently needed.

In recent years, liquid biopsies have emerged as a promising non-invasive alternative for the detection of multiple cancer types [23–27]. Such approaches use bodily fluids, for example blood, to detect specific disease-linked biomarkers, such as tumor circulating cells, proteins, or cell-free DNA. Similar efforts have emerged for PM, with a focus on the identification of circulating protein biomarkers. While studies have shown the potential of fibulin-3, mesothelin, and osteopontin, among others, their sensitivity and specificity for routine clinical use have not been confirmed yet [28–34]. At the same time, there is a limited number of studies focusing on RNA-based biomarkers for PM. Such biomarkers are particularly interesting, as they could shed more light on the transcriptional machinery involved in PM biology and potentially identify novel treatment targets.

Exosomes are small round-shaped endosomal vesicles, a subgroup of extracellular vesicles, secreted by all cells in an organism, including cancer cells [35,36]. They contain molecular cargo, with RNA, DNA, lipids and proteins, secreted by their donor cells. Several studies have already shown that cancer cell-secreted exosomes are shed into the bloodstream, suggesting them as a promising source of blood-based biomarkers [37,38]. Moreover, multiple studies have highlighted their role in cell-cell communication and reshaping phenotype of recipient cells [39,40]. For instance, cargo exchange between cells via extracellular vesicles has been confirmed for (1) RNAs between glioblastoma cancer cells [41], and glioblastoma and endothelial cells [42], (2) DNA sequences and retrotransposon elements between medulloblastoma and endothelial cells [43], (3) and for protein transfer between glioma cells [44]. Therefore, exosomes emerge as potential novel avenues for drug delivery systems, as well as mediators of drug resistance and disease progression [45,46]. We have previously shown that PM-derived extracellular vesicles carry RNA cargo enriched with genes associated with tumor progression, including long non-coding RNAs (LncRNAs), transcriptional factors, and marker genes of PM intratumoral transcriptional heterogeneity [47]. While our findings provide evidence for the diagnostic potential of secreted RNA cargo, the full extent of its clinical utility warrants further exploration.

In this work, we address this gap by comparing RNA cargo of exosomes secreted by PM (PM-exo) and non-PM (non-PM-exo) cells. By utilizing primary cell cultures established using pleural effusion and tissue samples from 11 PM and 6 non-PM patients, followed by exosome isolation and total RNA sequencing of RNA from exosomes, matching cells and tissues, we provide new evidence on the secretion of RNAs into PM-exosomes. We identified upregulated hallmarks of cancer associated with PM-exosome cargo, with high enrichment of genes linked to proliferation. We selected transcriptional factors, including *SETDB1*, *GATA2*, and *FOXM1*, which are potential regulators of up-secreted PM-exo genes, and whose cellular expression shows significant association with PM patient survival. Finally, we propose six secreted genes, including *GAS5*, *AL031666.3*, *RSLD1D1*, *AC103740.2*, *ADAM10*, and *AC020892.1*, as promising biomarkers for patient diagnosis, stratification, and prognosis assessment. Our comprehensive analysis provides novel insights into exosome-mediated PM biology, and proposes urgently needed biomarkers for non-invasive diagnostic approaches for PM detection.

## Results

### Molecular characterization of exosomes

In this work, we compared RNA cargo secreted by PM and non-PM cells, in order to identify RNA biomarkers for PM non-invasive diagnosis. To achieve this goal, we used diagnostic tissue samples and pleural effusion obtained from 11 PM and 6 non-PM, which were used to establish primary cells (Table 1, Figure 1A).

**Table 1.** Clinical characteristics of analyzed samples. Exo ID - ID of exosomal sample; Subtype - tumor histological classification; OS - overall survival time.

| Exo ID | Cell ID | Tissue ID | Group | Subtype | Age | Treatment | Stage | Gender | Asbestos exposure | OS |
| --- | --- | --- | --- | --- | --- | --- | --- | --- | --- | --- |
| RE41 | RCD41 | T41 | PM | biphasic | 68 | Cisplatin pemetrexed | III | male | possible | 26.35 |
| RE42 | RCD42 | T42 | PM | epithelioid | 50 | - | - | female | possible | 40.08 |
| RE43 | RCD43 | T43 | non-PM | - | 75 | - | - | male | no | - |
| RE45 | RCD45 | T45 | PM | epithelioid | 68 | - | IV | male | yes | 1.81 |
| RE46 | RCD46 | T46 | non-PM | - | - | - | - | female | - | - |
| RE47 | RCD47 | T47 | PM | biphasic | 56 | - | IV | male | yes | 3.47 |
| RE48 | RCD48 | T48 | PM | epithelioid | 69 | Carboplatin pemetrexed | - | male | yes | 13.07 |
| RE49 | RCD49 | - | non-PM | - | - | - | - | female | - | - |
| RE50 | RCD50 | T50 | PM | biphasic | 65 | - | III | male | yes | 11.47 |
| RE51 | RCD51 | T51 | PM | biphasic | 58 | Cisplatin pemetrexed | III | male | no | 15.97 |
| RE52 | RCD52 | T52 | non-PM | - | - | - | - | female | - | - |
| RE53 | RCD53 | T53 | PM | biphasic | 66 | Cisplatin carboplatin pemetrexed | II | male | yes | 12.78 |
| RE54 | RCD54 | T54 | PM | epithelioid | 90 | - | III | female | no | 4.80 |
| RE55 | RCD55 | T55 | non-PM | - | - | - | - | female | - | - |
| RE56 | RCD56 | T56 | PM | epithelioid | 64 | Cisplatin pemetrexed bevacizumab | II | male | yes | 14.98 |
| RE58 | RCD58 | - | non-PM | - | - | - | - | female | - | - |
| RE59 | RCD59 | T59 | PM | epithelioid | 84 | - | II | male | yes | 12.65 |

**Figure 1.**
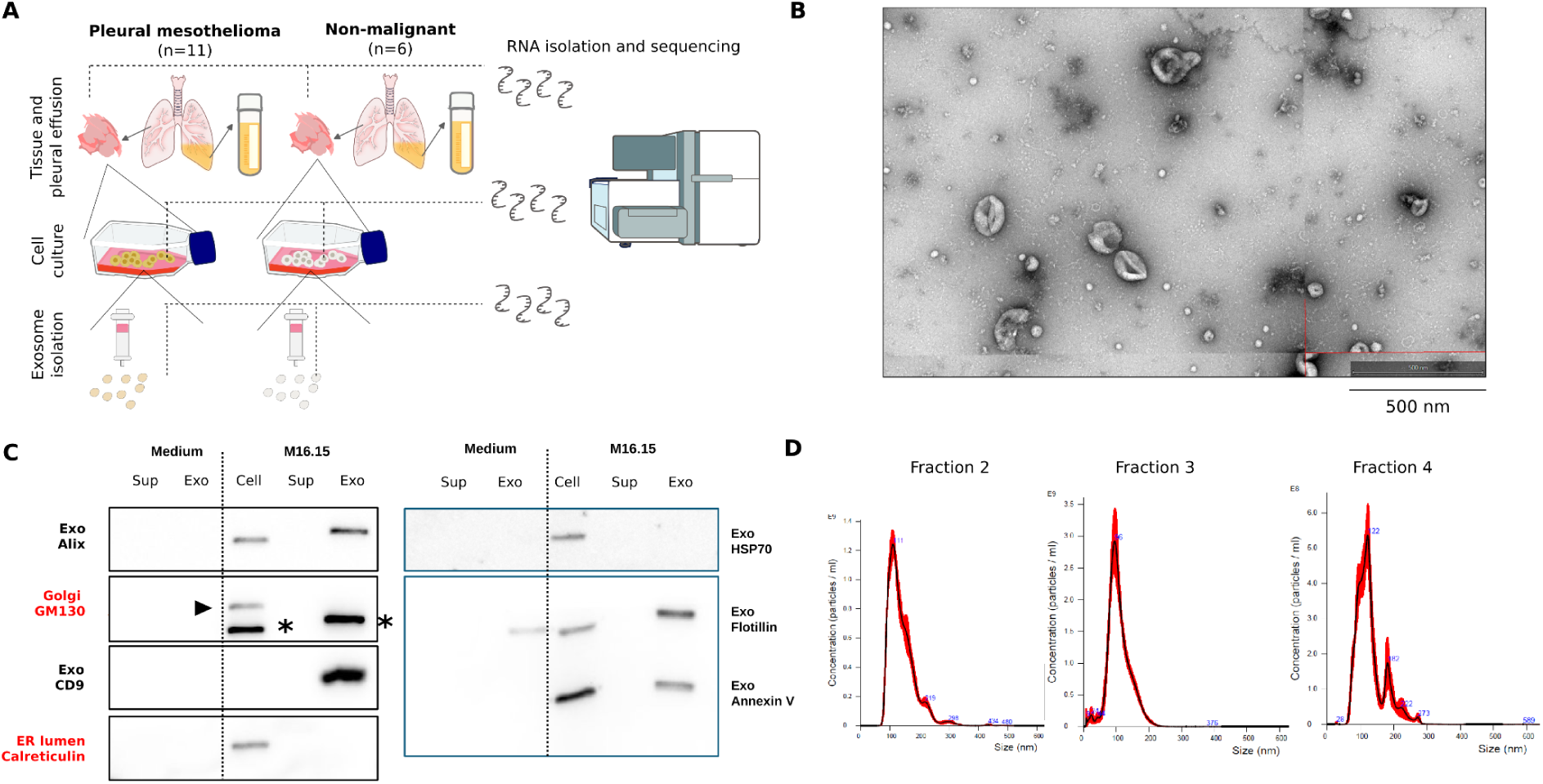
Exosome isolation and characterization. **A.** Diagram illustrating the workflow for screening exosomal RNA cargo. Tissue and pleural effusion samples from 11 PM and 6 non-PM patients were used. Tissue samples were used to establish primary cell culture. Supernatant of these cultures was used to isolate exosomes with qEV columns. RNA isolated from tissue, cells and exosomes was sequenced. **B.** Electron micrograph depicting exosomes isolated from M16.15 PM cell culture supernatant. **C.** Western blot analysis of exosomal and cellular organelle-specific proteins. Markers were tested in supernatant (Sup) and exosomes (Exo) isolated from cell culture medium and M16.15 cell culture. **\***remaining signal from Alix. **D.** Nanoparticle tracking analysis confirming enrichment of particles within the exosomal size range (mean diameter: 120 nm) in the collected eluate fractions from exosome isolation using qEV iZON column and M16.15 cell culture. PM - pleural mesothelioma; ER - endoplasmic reticulum.

The cellular origin of cultured cells was confirmed using immunohistochemical staining of epithelial and mesothelial markers, including pan-Cytokeratin, Podoplanin, Calretinin, and Wilms’ Tumor 1 (WT1) as described before [47]. To verify the agreement between genetic profiles of cultured tumor cells and matching tumors, we used Copy Number Arrays, in particular comparing the regions of *BAP1* and *MTAP*/*CDKN2A* genes, as those are often altered in PM. We found high agreement between cultured cells and the original tumors (Additional File 1). Comparison of transcriptomic profiles of these cells revealed two clusters, corresponding to PM and non-PM cells (Suppl. Figure S1A). To better characterize transcriptomic profiles of PM cells, we used previously published *S-* and *E-*component marker genes [19], and scored their expression in cells. Except for four cells, the scores corresponded to tumor histological type (Suppl. Figure S1B-C).

In the next steps, we isolated exosomes using cell culture supernatant and qEV columns (iZON), a size-exclusion chromatography technique. Purity of isolated vesicles was confirmed with Nanoparticle Tracking Analysis (NTA), by measuring the size distribution of vesicles obtained in individual qEV fractions (Figure 1D, Suppl. Figure S2). Given the highest enrichment of vesicles in the expected exosome size in fractions 2-4, vesicles from these fractions were pooled together and used for downstream analysis. Purity of exosomes was also verified by western blot analysis of exosome-specific proteins, i.e. Alix (cytosolic protein), CD9 (transmembrane protein), HSP70 (cytosolic protein), Flotillin (cytosolic protein with lipid or membrane binding capacity), and Annexin V (a phospholipid-binding protein) (Figure 1C). Their expression was tested separately in cells, supernatants, and exosomes. In addition, we tested the expression of markers associated with the Golgi apparatus matrix (GM130) and the endoplasmic reticulum lumen (Calreticulin). As expected, exosome-specific markers were expressed in cells and exosomes, while GM130 and Calreticulin were observed only in cells (Figure 1C), confirming purity of isolated exosomes. In addition, Electron Microscopy images confirmed the expected size and morphology of isolated exosomes (Figure 1B).

### Transcriptomic cargo of exosomes and matching cells

In the next steps, we isolated RNA from exosomes, matching cells and tissues, and followed by RNA-sequencing (Methods). After data preprocessing, reads were mapped to the human reference genome GRCh38 and gene expression was quantified.

In total, we found 34,535 and 31,554 genes expressed across all PM-exo and non-PM-exo samples, respectively. The majority of genes in PM-exos were protein coding genes (48%), long noncoding RNA (LncRNA, 33%), and pseudogenes (18%) (Figure 2A, Suppl. Table S1). In case of non-PM exosomes, we observed a similar trend of gene biotype distribution, with a slightly higher proportion of protein coding genes (Figure 2A, Suppl. Table S1).

**Figure 2.**
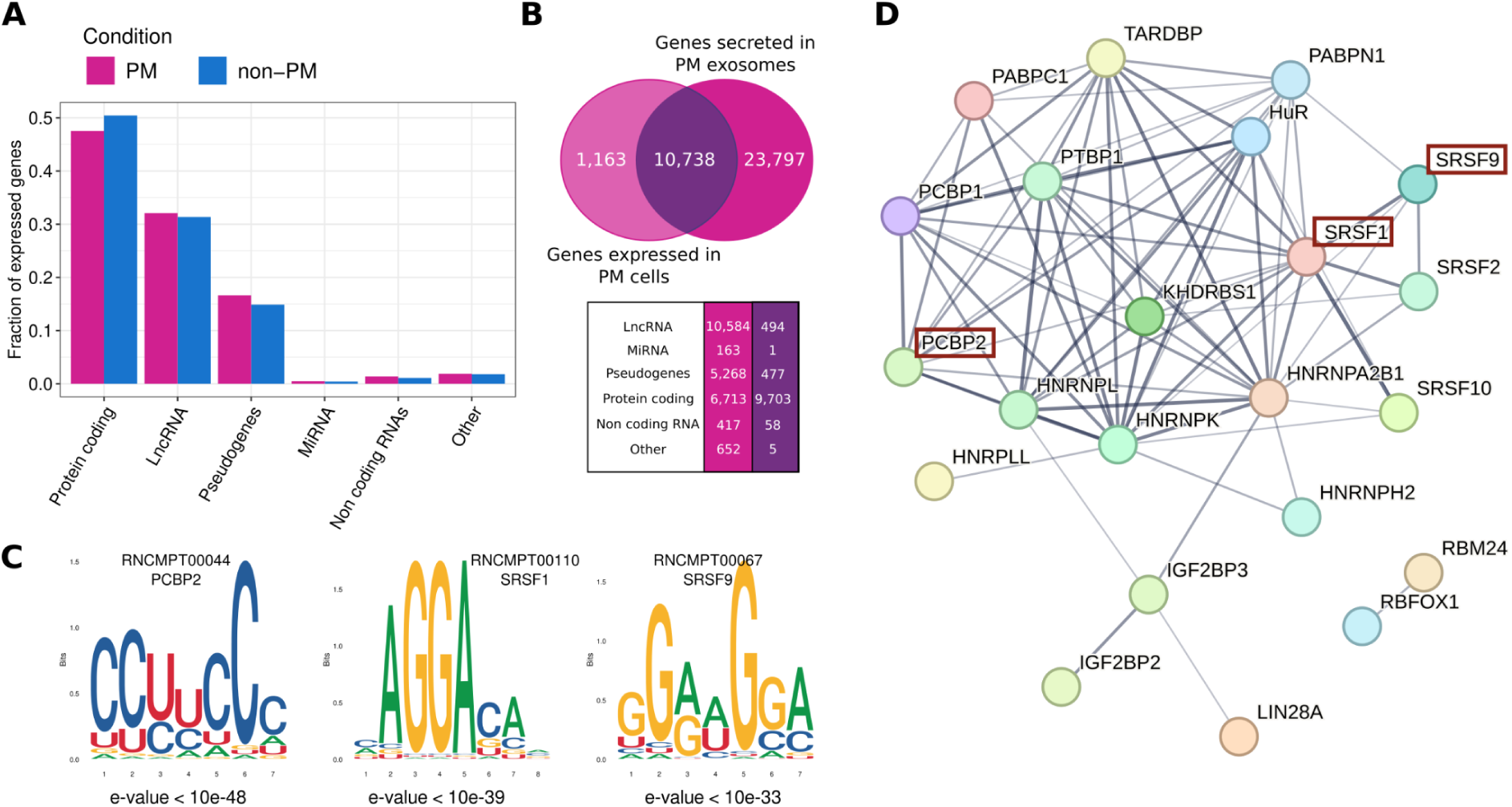
RNA cargo and regulatory motifs in PM-exosomes. **A**. Distribution of biotypes among genes expressed across all PM and non-PM exosomes. **B.** Comparison between PM-exosomal RNA cargo and genes expressed in matching PM cells. Absolute number of genes shared between exosomes and cells, or present only in exosomes, is reported in the table. **C.** Sequence logo of the three most enriched binding motifs of the PM-exo up-secreted genes. **D**. Protein-protein interactions of the RNA-binding proteins associated with significantly enriched motifs in PM-exo transcripts. Red boxes highlight the three proteins from **C**.

To gain better understanding on the link between exosomal cargo and RNAs expressed in matching cells, we analyzed distribution of gene biotypes present in cells. While in total 11,901 genes were expressed across all PM cells, over 80% of them represented protein coding genes, with significantly lower fraction of LncRNA and pseudogenes compared to exosomal samples (Suppl. Figure S3, Suppl. Table S2). Similar trend was observed in non-PM cells (Suppl. Figure S3, Suppl. Table S2).

We found 10,738 genes shared across PM-cells and PM-exos, with 1,163 genes exclusively expressed in PM-cells, and over 23,797 genes exclusively present in PM-exo (Figure 2B). As expected, the majority of the RNAs specific to PM-exo were non-coding RNAs and pseudogenes. Similar results were observed in non-PM samples (Suppl. Fig. S4), confirming higher enrichment of non-coding RNAs in exosomes compared to matching cells.

To identify potential differences in RNA cargo of microvesicles, another group of extracellular vesicles (EVs), and exosomes, we used our previous results on the RNA cargo of mixed population of EVs isolated from the same PM cells as those used in current study (Suppl. Table S3) [47]. In total, we found 11,405 genes differentially expressed between EVs and exosomes, with 817 genes upregulated and 10,588 downregulated in exosomes (log2 fold change threshold 1 and -1, p-value < 0.05, Suppl. Fig. S5). Interestingly, the downregulated genes in exosomes were mostly protein-coding genes (9,675 genes; Suppl. Table S4).

### RNA-binding proteins involved in PM-exosome cargo sorting

Given the observed varied distribution of detected genes in matching EVs, exosomes and cells, we investigated the potential mechanism of selective cargo sorting into exosomes. Specifically, we used Analysis of Motif Enrichment (AME) from the MEME suit [48], to analyze the enriched RNA-binding motifs in PM-exo genes. We found strong enrichment of 42 motifs (e-value < 0.05) associated with several RNA-binding proteins (Suppl. Table S4). Among top significant hits, we found CU-rich motifs bound by Pcbp2, and GGAG-rich motifs, bound by Srsf1 and Srsf9 (Figure 2C). Srsf1 has been previously shown to mediate selective sorting of microRNA into exosomes in pancreatic and breast cancer cell lines [49], while Pcbp2 has been associated with higher microRNA secretion via EVs, leading to EGFR-driven angiogenesis [50]. Both Srsf1 and Srsf9 binding was shown to promote Beta-catenin accumulation in cancer cells, which promoted Wnt signalling [51].

Additionally, we used the STRING database [52], which collects experimentally verified protein-protein interactions, to identify additional proteins involved in the PM-exo cargo sorting and interacting with identified RNA-binding proteins. We identified 13 additional proteins highly interacting with Pcbp2, Srsf1 and Srsf9, including hnRNPA2B1, previously shown to regulate microRNA loading into exosomes [53] (Figure 2D).

### Transcriptomic comparison of PM and non-PM-exosomes

Given the growing evidence of exosomes as a source for disease biomarkers, we compared the transcriptomic cargo of exosomes secreted by PM and non-PM-cells. Overall, 3,204 and 1,103 genes were up- and down-expressed in PM-exo compared to non-PM-exo samples (Figure 3A, p-value < 0.05, Suppl. Table S5). Among top enriched genes, we found four *S-*component and three *E-*component genes (Suppl. Figure S6A), along with 20 transcriptional factors (Suppl. Figure S6C), 16 enriched microRNAs (Suppl. Figure S6B) and 720 LncRNAs (Suppl. Figure S6D, Suppl. Table S5). Interestingly, among the up-secreted LncRNAs we found *GAS5* and *POT1-AS1*, previously identified as potential blood-based biomarkers for epithelioid PM [54]. In addition, *ABCA5*, *ABCA10* and *ABCA9-AS1*, which represent ABC-transporter genes (Suppl. Fig. S6E), previously associated with the multidrug resistance phenotype [55], were among the up-regulated genes.

**Figure 3.**
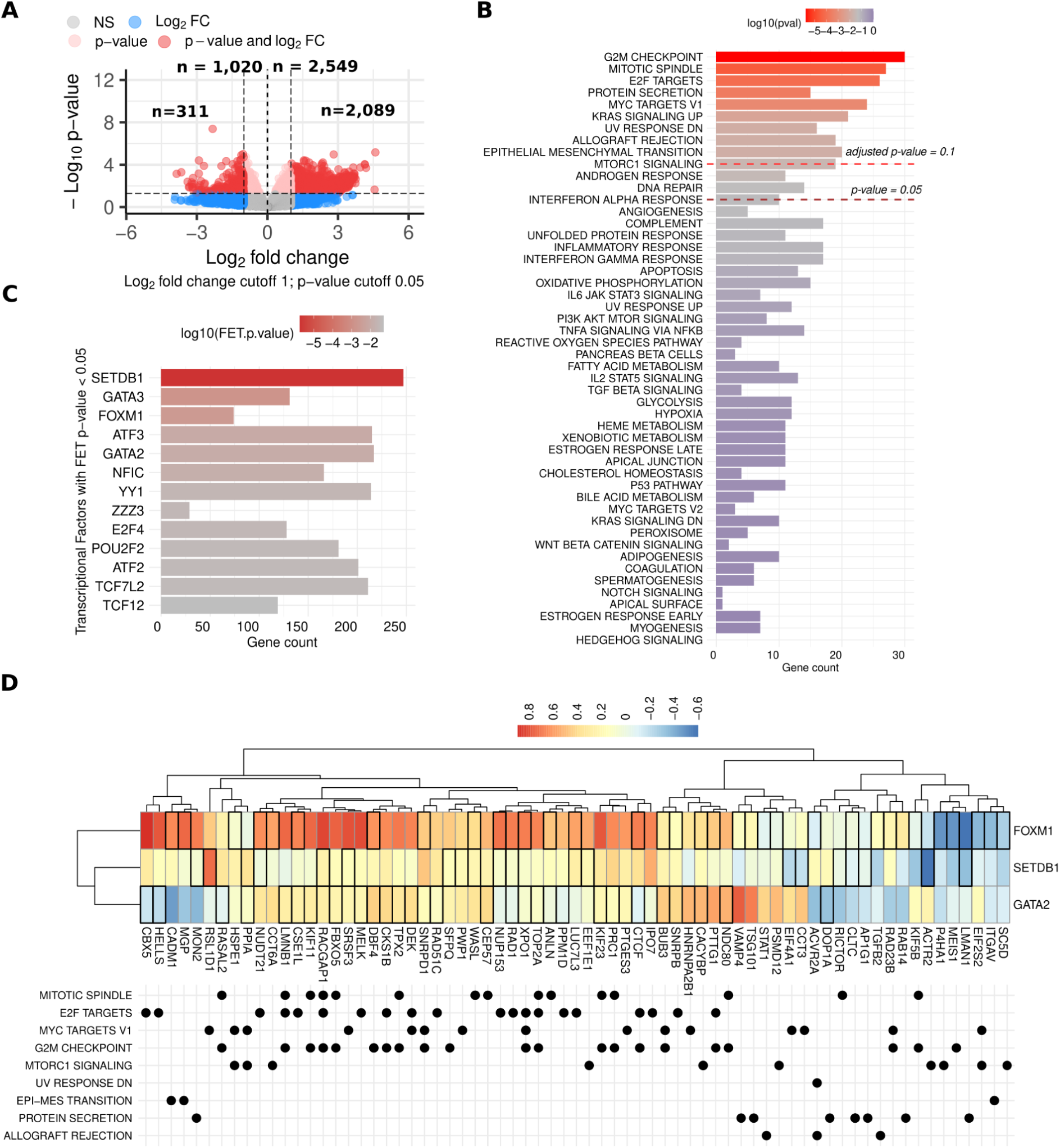
Functional and regulatory landscape of PM-exosomal cargo. **A.** Differential gene expression between PM and non-PM exosomal cargo. **B.** Hallmarks of cancer in which PM-exo up-regulated genes are involved. **C.** Top 10 transcriptional factors with the strongest enrichment of PM-exo up-secreted genes. **D.** Correlation between expression of the top three most enriched transcriptional factors and their target genes expression in PM cells. Color scale represents Pearson’s correlation coefficient. For each target gene, hallmarks of cancer in which it is involved is marked below the heatmap.

### Hallmarks of cancer enriched in PM-exosomes

To understand processes in which PM-exosome enriched genes are involved, we used gene set enrichment analysis to verify their association with hallmarks of cancer from MsigDB. We found significant enrichment of 13 hallmarks (p-value < 0.05), with *G2M Checkpoint*, *Mitotic spindle formation*, and *E2F targets* among the top enriched pathways (Figure 3B).

In the next steps, we scored exosomal expression of the genes involved in the top enriched hallmarks, as well as up-secreted *S-* and *E-*component genes, to assess the link between exosomal activity of these pathways and PM patients’ survival. For each score, we used a multivariate Cox regression model along with tumor histological subtype and treatment information (Table 1). Higher scores of seven pathways were associated with worse patient prognosis, with the highest hazard linked with *UV response down* set (Suppl. Figure S7). Interestingly, *Epithelial-Mesenchymal Transition* (EMT)-related score was associated with better patient prognosis, contrary to the growing evidence on the link between EMT and cancer progression [56]. In parallel, we scored the expression of the same genes in PM cells, and repeated Cox regression analysis to assess the link between cellular activity of these pathways and patient survival (Suppl. Fig. S9). Four sets, i.e. *MYC targets*, *MTORC1 signaling*, *E-component*, and *E2F targets*, showed association with higher hazard, while three sets, i.e. *Allograft rejection*, *KRAS signaling up*, and *EMT*, were linked with lower hazard, consistent with results found for exosomes. Interestingly, higher levels of *UV response down*-related genes in PM cells were linked with better survival, contrary to their link with higher hazard based on exosomal expression (Suppl. Figure S7-S8).

### Transcriptional regulators of PM-exosomal RNA cargo

To identify potential transcriptional factors (TF) regulating expression of PM-exo enriched genes, we used ChEA3 [57], which provided enrichment scores of TFs with experimentally verified binding sites to promoter regions of PM-exo enriched genes. We obtained 13 TFs with significant enrichment (FET p-value < 0.05, Suppl. Table S7), with *SETDB1*, *GATA3* and *FOXM1* among the top enriched factors, targeting respectively 248, 127 and 75 genes expressed in PM-exos (Figure 3C, Suppl. Table S7). From the set of 13 enriched TF, cellular expression of *SETDB1*, *FOXM1* and *GATA2* was significantly linked to patient survival (Cox multivariate regression analysis with subtype and treatment information, p-value < 0.05, Suppl. Figure S9). *GATA2* was predominantly positively correlated with expression of its targets, mostly involved in proliferation, i.e. *MYC targets V1*, *E2F targets*, and *G2M checkpoint* (Figure 3D). *FOXM1* was the only TF regulating genes involved in EMT, with strong positive correlation, while both *FOXM1* and *SETDB1* negative regulated genes involved in *mTORC signaling* (Figure 3D).

### Technical validation of candidate RNA biomarkers using RT-qPCR

Next, we selected 30 candidate genes for technical validation using technical replicates of PM and non-PM cell cultures whose secreted exosomes were profiled with RNA-sequencing (Suppl. Figure S10). We repeated exosome isolation from the technical replicates of primary cells, isolated exosomal RNA, and used quantitative reverse transcription polymerase chain reaction (RT-qPCR) to validate the enrichment of selected genes in PM- vs. non-PM-exo. The analysis confirmed significantly higher expression of 7 LncRNA and 14 protein coding genes in PM compared to non-PM exosomes (Figure 4A-B, Suppl. Table S6, S8; Mann-Whitney p-value < 0.1).

**Figure 4.**
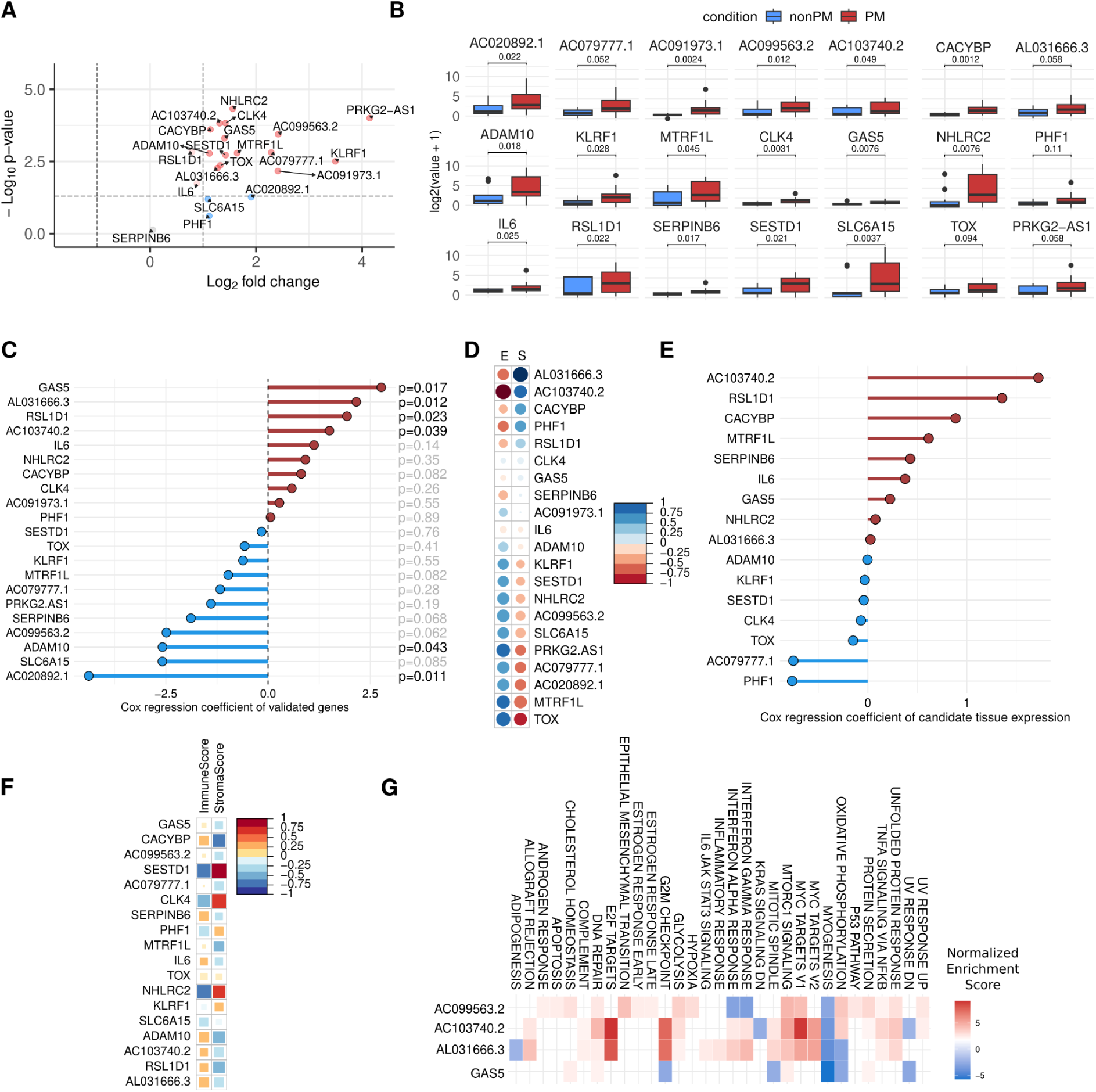
Exosomal candidate genes for PM detection. **A.** Volcano plot generated using sequencing data of qPCR-validated candidate genes. **B**. RT-qPCR validation of PM-exo candidate genes in technical replicates of exosomes. Significance was computed using Mann-Whitney U-test. **C.** Cox survival analysis using exosomal expression of validated candidate genes (multivariate models with histological type and treatment). **D.** Correlation between exosomal expression of candidate genes and *S-/E-*compartment scores of matching PM cells. **E.** Cox survival analysis of candidate genes in PM tissue expression data. **F.** Correlation between PM tissue expression of candidate genes and xCell Immune and Stromal scores. **G.** Hallmarks of cancer enriched with predicted targets of validated long non-coding RNA genes.

Six of the validated genes showed significant association with patient survival (Cox regression p-value < 0.05, models including histological subtype and treatment information, Figure 4C). The validated genes in general had higher expression in epithelioid tumors compared to biphasic ones (Suppl. Fig. S11), with five candidates, *AL031666.3*, *AC103740.2*, *CACYBP*, *PHF1* and *RSLD1,* were positively correlated with *E-*scores of matching cells. Conversely, ten genes, *KLRD1*, *SESTD1*, *NHLRC2*, *AC099563.2*, *SLC6A15*, *PRKG2-AS1*, *AC079777.1*, *AC020892.1* and *TOX*, were positively correlated with *S-*component scores (Figure 4D).

### Potential role of PM-exosomal RNA cargo in the tumor microenvironment

In the next steps, we used RNA-sequencing data from exosome-matching tissue samples, to assess the clinical and functional role of validated PM-exo genes in the tumor microenvironment (TME). The expression of *CACYBP*, *AC103740.2*, and *RSL1D1* was predictive of patient survival, in both univariate and multivariate models with subtype information (Figure 4E, Suppl. Table S9). Survival analysis using TCGA mesothelioma data confirmed results for the two first genes, and revealed three additional genes with significant association (Cox univariate p-value < 0.05, Suppl. Fig. S12).

In the context of TME composition of PM tissues, we found an association between tissue expression of several candidate genes with either immune or stromal scores, reflecting the infiltration of the respective cell types. Specifically, *SESTD1*, *CLK4* and *NHLRC2* showed the strongest correlation with stroma scores, with *CACYBP*, *SERPINB6*, *IL6*, *ADAM10*, *AC103740*.2, *RSL1D1* and *AL031666*.3 positively linked to immune scores (Figure 4F). We observed similar patterns of association in TCGA mesothelioma data (Suppl. Fig. S13). Interestingly, several of the genes, in particular *AC103740.2* and *TOX*, were also positively correlated with PD-L1 protein expression as well as CD274 gene expression in TCGA tissue samples, suggesting they could potentially be further explored for immunotherapy patient stratification (Suppl. Fig. S14). In addition, we found four genes with significant differences in gene expression across tumor subtypes (Suppl. Table S10, Figure S19), as well as higher expression of *AC103740.2* associated with better response to chemotherapy in TCGA mesothelioma samples (Suppl. Fig. S15).

Finally, using PM tissue gene expression, we identified potential protein coding target genes of four validated LncRNA biomarkers, which were enriched with over 30 different hallmarks of cancer (adjusted p-value < 0.01, Figure 4F). The target genes presented the strongest enrichment of *E2F targets*, *G2M checkpoint*, and *MYC targets V1* pathways, and negative enrichment of *Myogenesis, Oxidative phosphorylation*, and *UV response down*, which suggest they could be potential co-activators and repressors of these respective processes in PM-exosome-recipient cells.

## Discussion

Exosomes, previously explored in other cancer types and highlighted as important mediators of tumor progression and cellular cross-talk, have not yet been fully investigated in PM. Our analysis, which integrates transcriptomic data from exosomes secreted by PM and non-malignant cells complemented with RNA-sequencing data from matching cells and tissues, allowed us to explore exosome-mediated PM biology at multiple levels.

First, as previously reported, we observed a higher proportion of non-coding RNAs in exosomes compared to matching cells, emphasizing their role as potential mediators of post-transcriptional regulation in recipient cells [58–60]. When comparing genes secreted into PM exosomes and mixed population of extracellular vesicles, we found higher expression of protein coding genes in the mixed population, suggesting that exosomal and microvesicle cargo may be sorted using different principles, therefore may have distinct functions in recipient cells.

Second, we identified several RNA-binding proteins that may be involved in selective sorting of RNAs into exosomes. Among the most associated RNA-binding proteins, we found Pcbp2, Srsf1 and Srsf9, previously linked with microRNA binding and secretion in cancer cells [49][50][51]. Since exosomal RNA cargo can contain molecules capable of inducing pro-oncogenic changes in recipient cells, identifying proteins involved in its sorting could be crucial for future targeting of pathways regulating their secretion.

Third, the comparison between exosomes secreted by PM and non-malignant mesothelial cells, allowed us to identify over 2,000 genes with higher expression in PM, including transcriptional factors and non-coding RNAs. We confirmed 21 candidate biomarkers, including long noncoding RNAs and protein-coding genes, such as *GAS5*, *ADAM10*, *AL031666.3*, *RSLD1D1*, *AC103740.2*, and *AC020892.1,* using an orthogonal experimental approach. Importantly, exosomal expression of these genes was significantly linked with patient survival in our cohort, and several of them were correlated with scores characterizing PM-specific intratumor transcriptional heterogeneity.

The exact role of exosomal cargo in cancer has not been fully explored yet, however, there are two main hypotheses. One suggests that the cargo consists mostly of cellular by-products destined for removal, while the other suggests that RNAs are selectively sorted into exosomes for intercellular communication. To explore these hypotheses, we analysed enriched pathways in which PM-exo genes are involved, as well as their association with patient survival. Overall, PM-exo cargo was enriched with hallmarks of proliferation, suggesting that exosomes could potentially transfer proliferation-linked phenotypes into their recipient cells. While we have not tested the functional role of the cargo in recipient cells, the already published evidence supports its potential role in modulating phenotype of recipient cells [39,61].

Finally, we explored the potential effects of the validated candidates in recipient cells in TME. We identified several genes that could be targeted by enriched LncRNAs in tissue samples, and processes in which they are involved. The analysis showed that *AC103740.2*, *AL031666.3* and *AC099563.2* may promote proliferation, while *GAS5* could be linked with decreased activity of oxidative phosphorylation, both processes important for tumor progression. Moreover, *AC103740.2* expression in tissue was associated with higher immune TME scores, suggesting it might have higher expression in immune-rich tumors. This was consistently observed in both our cohort and TCGA mesothelioma data. In addition, its expression in TCGA data was associated with PD-L1 expression, suggesting it could potentially be used as a biomarker for immunotherapy patient stratification.

Our study has several limitations. First, our analysis was limited to tumors with epithelioid and biphasic subtypes, potentially limiting the power of identifying biomarkers of sarcomatoid subtype. However, we observed that the molecular classification of tumours did not directly correspond to the enrichment of *S-* and *E-*scores of matching PM cells, with our results showing that some cells originating from epithelioid tumors presented high expression of the sarcomatoid-like component, and vice versa. Therefore, we believe that the identified biomarkers are universal for PM detection across histological subtypes. Second, our analysis on the proteins involved in cargo sorting, transcriptional factors regulating PM-exo genes, and functional effects of PM-exosome on recipient cells, should be verified experimentally to validate their involvement in these processes. Third, extensive validation of the identified biomarkers in large cohorts of plasma samples is necessary to confirm their diagnostic and clinical potential.

Despite these limitations, our work presents new insights on exosomes secreted in pleural mesothelioma. It is the first work to characterize their RNA composition and evaluate their clinical significance in the context of matching cells and tissue samples. Our findings provide new evidence on the role of exosomal RNA in PM biology and highlight their potential as a source for non-invasive circulating biomarkers.

## Materials and methods

### Cell cultures

Primary cell cultures of 9 pleural mesothelioma and 6 non-malignant mesothelial cells were established using treatment-naive pleural effusion samples of patients undergoing diagnosis at the Department of Thoracic Surgery, University Hospital Zurich (Table 1). Two cell cultures were isolated from PM tumor tissues as previously described, SDM90 and SDM134 [62]. Pleural effusion samples were processed as described previously (ref our paper)and cell cultures were maintained at 37 °C and 5% CO_2_ and verified to be free from mycoplasmas. To identify the identity of the cells, we performed immunohistochemical staining of protein markers as described previously [47] and copy number variation array. Four PM and 2 non-PM cell lines were validated previously [47]. The verification of additional cell lines used in this study is provided in the Additional File 1.

### Exosome isolation

Cells were seeded in the complete medium in 10 of 150 cm^2^ non-coated cell culture dishes until reaching 50–70% confluency. Cells were then washed three times with 15 ml DPBS to clean exosome contamination in the FBS. After that, 20 ml of the culture media, RPMI medium ATCC modification (GibcoTM, A1049101, Life Technology Europe BV, Zug, Switzerland) containing 2mM glutamine (Sigma-Aldrich Buchs, Switzerland), human Epidermal Growth Factor (20 ng/ml, hEGF) and 8% exosome depleted FBS (Life technology A27208-03) was added to each dish. After 72 h incubation, we collected the all 20 ml supernatant and cells were washed again with 10 ml DPBS to collect the remaining medium. The total volume of 300 ml supernatant plus DPBS was collected from each cell culture. The medium was centrifuged at 400 g for 5 min at 4°C to sediment cells. The supernatant was subjected to the next centrifugation again at 3,000 g for 15 min at 4°C to remove large debris. Afterwards, the supernatant was filtered through a 0.22 µm vacuum filtration device (TPP, 99250). The supernatant was further concentrated to a total volume of 4 ml using Centricon Plus-70 PL-100 100k (Millipore UFC710008). The concentrate was again centrifuged at 10,000 g for 10 minutes at RT to remove debris. Afterwards, we performed exosome isolation using iZON qEV1/35 nm (iZON Science) following manufacturer’s guidelines using loading volume of 1 ml. We collected and pooled fraction 2-4 as these are enriched with exosomes (supplementary figure X). The exosome isolation was repeated 4 times using the same iZON column with in between washing with 1 column volume (14 ml) of DPBS. Exosomes solution was either directly used for NTA and electron microscopy. For RNA and protein isolation, we added 800 ul Nanotrap® EV Particles (RNA concentration kit, iZON Science) in 8 ml exosome eluate, incubated and rotated at room temperature for 1 hour. After that, the samples were centrifuged at 16,800 g for 10 minutes and the supernatant was discarded. The remaining Nanotrap® pellets bound to exosomes were snap frozen and stored at -80°C until further analysis.

In addition to exosomes, we also collected the cells (from 1 dish) by trypsinization and counted the number of cells and verified the cell viability of >80% for all samples. Cells were washed twice in DPBS and snapped frozen for RNA isolation.

### Western blot analysis

Nanotrap® pellets bound to exosomes or cells were lysed with cell lysis RIPA buffer (50 mM Tris HCl pH. 8, 150 mM NaCl, 1% TritonX-100, 0.5% sodium deoxycholate, 0.1% SDS) containing protease/phosphatase inhibitor cocktail (Cell Signaling, #5872, Danvers, MA, USA). For supernatant we used 4 ml of supernatant and concentrated using Amicon Ultra 4 ml 3K Preparation of cell extract and western blot, we performed as we previously described [47]. We used the primary and secondary antibodies from exosomal marker antibody sampler Kit (Cell Signaling, Danvers, MA, USA, #74220) to detect the expression of exosomal markers and markers for possible contamination from released organelles including Golgi and Endoplasmic reticulum.

### Copy Number Analysis

We isolated DNA from primary cell cultures using Qiagen DNeasy Blood & Tissue mini Kits (69504, Qiagen Hombrechtikon Switzerland). DNA from tumor tissues were isolated by Qiagen FFPE DNA isolation kit (56544, Qiagen). PM primary cells and matched tissue samples were characterized using Affymetrix OncoScan Microarray according to the instruction manual (Applied Biosystems, 902695). Obtained CEL files were analyzed using Analysis Power Tools (Thermo Fisher, version 2.4.0) and rCGH R package. The results of the analysis are presented in Additional File 1.

### Nanoparticle Tracking Analysis

Nanoparticle Tracking Analysis was performed using NanoSight (NS300-Malvern).

### Electron Microscopy

5 µl of elute from the iZON qEVoriginal column was applied to a glow-discharged grid of 300 mesh copper and incubated for 1 minute. Excess liquid was removed and then stained with, with 2% uranyl acetate for 1 min. The slide was dried and subjected to imaging by transmission electron microscopy.

### RNA isolation

We used *mir*Vana PARIS Kit (Ambion #AM1556) for RNA isolation. We added 625 ul ice cold cell disruption buffer to the cell pellet, after vortexing we added equal volume of Phenol-Chloroform to the sample and proceeded with totalRNA isolation according to the instruction manual. Nanotrap® pellets bound to exosomes were resuspended in 400 ul ice cold cell disruption buffer. After vigorous vortexing, the sample was centrifuged at 16,800 g, 4°C for 10 minutes. The supernatant was collected for RNA isolation and the equal volume of Phenol-Chloroform was added followed by RNA isolation. For tumor tissues, we used tissue samples frozen in optimal cutting temperature (OCT) medium. We performed tissue sectioning (10 um) with a microtome. Tissue sections were lysed with 625 ul ice cold cell disruption buffer and proceed as for RNA isolation from cells. From 2 samples (SDM90 and SDM76), the RNA were previously isolated with the RNeasy kit (Qiagen). A total of 85 ul RNA eluate was obtained from the *mir*Vana PARIS kit. RNA concentration was measured by Qubit™ RNA High Sensitivity assay Kit (Invitrogen, Q32852). RNA concentration from exosome samples were below the detection limit despite the use of the high sensitivity kit. The RNA from cells and tissues were treated with DNAse prior to library preparation using TURBO DNA-free™ Kit (Ambion AM1907) according to manufacturer’s instructions. RIN number of RNA was measured by bioanalyzer device using Eukaryote Total RNA Pico kit ensuring RIN>7 for all tissue samples with the exception of M21.04, M16.25 and M19.36.

### Preparation of libraries

As starting material, we used 5 ng of DNase treated RNA extracted from cells or tissues in a total volume of 7 µl. For exosomal RNA, we used a total of 7 µl of RNA suspension as the starting material from all samples, because the RNA concentration was below the detection limit. For all samples, we performed 6 minutes of RNA fragmentation. We used SMART-Seq Stranded Kit (Takara, 634444), together with the SMARTer RNA Unique Dual Index Kit – 96U Set A, (634452) for the indexes. We used 5 cycles for PCR-1 step and 13 cycles for PCR-2 step. One final clean up step was applied. Library size was verified using bioanalyzer. Library preparation was conducted for cell culture medium (negative control), where no library amplification was detected.

### RNA-sequencing data analysis

All libraries were pooled and sequenced using a Novaseq6000 SP(100) in one S4 lane, to obtain paired-end 150 bp reads. Data was sequenced at the Genomics Facility Basel. Quality of obtained reads were accessed using FastQC (https://www.bioinformatics.babraham.ac.uk/projects/fastqc/, version 0.11.9). Reads were processed using fastp [63] to remove adapter sequences and perform quality-based read trimming. Processed reads were mapped to the human reference genome GRCh38.p13 using STAR (https://github.com/alexdobin/STAR; version 2.7.5c). STAR-inferred gene counts were used for downstream analysis. Annotation of transcriptional factors was obtained from the FANTOM5 database (https://fantom.gsc.riken.jp/5/). *S*- and *E-*component marker genes were obtained from [19]. For UMAP visualisation of cells, calculated using the UMAP R package [64], data was normalized using Variance Stabilized Transformation (vst) function from the DESeq2 R package [65].

### Enrichment of RNA-binding proteins

Sequences of the longest transcripts of the PM-exosome expressed genes were obtained from the gencode GTF file (version 35). Analysis of Motif Enrichment from MEME suite [48] was used along with the Ray2013 database of RNA-binding proteins, to identify proteins whose binding motifs are enriched in the PM-exo transcripts, with transcripts expressed in matching PM-exo cells as as background set. Identified proteins whose motifs had significant association (p-value < 0.05) were provided to the STRING database [52] to identify interacting proteins based on experimental data and medium interaction confidence (score > 0.4).

### Differential gene expression analysis

Genes upregulated in PM-exosomes compared to non-PM-exosomes were identified using differential gene expression analysis with the DESeq2 R package [65]. For the biotype analysis of expressed genes, the gencode GTF file (version 35) was used.

### Gene set enrichment analysis

Gene set enrichment of the upregulated PM-exosome genes was calculated using hallmarks of cancer from MsigDB (https://www.gsea-msigdb.org/gsea/msigdb/) and hypergeometric test from the fgsea R package (https://bioconductor.org/packages/release/bioc/html/fgsea.html). *S-* and *E-*compartment scores of PM cells, tissue, TCGA expression, as well as scores of selected enriched pathways in exosomes, were calculated using the ssgsea [66] method from the GSVA R package.

### Analysis of Transcriptional Factors

Chea3 (https://maayanlab.cloud/chea3/) was used to identify transcriptional factors whose experimentally validated targets were enriched in the PM-exosome enriched genes. For evaluation of regulation effect on target genes, we summed normalized counts from exosomes and matching cells, as total expressed genes, and correlated these values of the PM-exosome enriched genes with the transcriptional factors identified with Chea3.

### Survival analysis

Cox regression analysis implemented in the survival R package was used to assess association with survival. Multivariate models were trained using respective genes or scores calculated for cells, exosome or tissue samples, along with tumor subtype and treatment, provided in Table 1. For TCGA mesothelioma data, tumor subtype information was used in multivariate models.

### RT-qPCR analysis

We used Prelude One-Step PreAmp Master Mix (Takara Bio, 638553) optimized for unbiased co-amplification to pre-amplify the cDNA generated from exosomal RNA to increase the detection sensitivity. For gene specific preamplification, the final concentration of each primer was 50 nM in a total volume of 25 ul reaction. As starting material, 5 µl of RNA solution from the miRVANA PARIS kit eluate was used. The preamplification was performed using the following cycling condition for each primer mix: 42°C 10 minutes, 95°C 2 minutes, 14 cycles of 95°C10 second and 60°C for 4 minutes. Using this reaction setting, high efficacy of pre amplification was observed (Ct Preamp- Ct without Preamp > 10 cycles). For qPCR, we diluted the preamplification reaction to at least 1:2.2 dilution with ultrapure water. A positive control (RNA isolated from cells) and no template control (ultrapure water) were included in every set of preamplification. Three µl of the diluted sample was used for quantitative real-time PCR (qPCR) in at least technical duplicates, using KAPA SYBR FAST qPCR master mix with low ROX (Kapa Biosystems, KK4621 Merck, Buchs, Switzerland) in a 10 µl reaction volume containing 200 nM of each primer, using the same instrument and setting as previously described. No template control for qPCR reaction was conducted. cDNA from no-preamplification was run on every plate for the comparison of melt curve analysis. We assumed a reaction efficiency of 100%. To normalise the expression data we selected genes that showed no difference in the expression between PM-exo and non-PM-exo, and gene candidates previously proposed as reliable and stable housekeeping genes for exosomes [67]. Among the selected genes, we identified OR2V2, OST4 and SNPRG that showed no significant difference in Ct between PM and non-PM exosomes (Suppl. Fig. S20). In addition, we observed a tendency of negative correlation between cell number and Ct value of these genes (Supp. Fig. S21). Gene expression was quantified using **2**^-ΔΔCT^ method normalized with average Ct of the reference genes (OST4, SNRPG, and OR2V2) using mean ΔCt of non-PM exosome samples as reference value. OR2V2 was included in all the primer mix to ensure the consistent preamplification efficiency of every preamplification. Primers used are listed in Suppl. Table S11.

### Comparison between matched EV and exosome RNA cargo

We used extracellular vesicle RNA-seq data from [47] to compare RNAs present in EVs and exosomes secreted from matching cells. ID matching of the EV and exosome samples can be found in Suppl. Table S3. For this comparison, the expression of the exosome samples was calculated using Kallisto (https://pachterlab.github.io/kallisto/download.html; version 0.46.1) as described before [47]. Then, EV and exosome expression was batch-normalized using the Qsmooth R package (https://bioconductor.org/packages/release/bioc/html/qsmooth.html). Differential gene expression was analyzed using DESeq2 R package [65].

### TCGA mesothelioma data analysis

STAR counted and FPKM-UQ normalized RNA-seq data from GDC TCGA mesothelioma dataset was downloaded from UCSC Xena browser (https://xenabrowser.net/datapages/). Log2 transformation was reversed, and used for downstream analyses. For clinical association analysis, clinical and treatment information linked to the dataset was obtained using TCGAbiolinks R package following the instructions from user manual [64,68]. The significance of associations was tested using Kruskal-Wallis test. The normalized RPPA expression of PDL1 for correlation analysis with gene expression of PM-exo genes, was downloaded from UCSC Xena browser (https://xenabrowser.net/datapages/).

### Tumor Microenvironment Scores

The scores representing infiltration of immune and stromal cell types of TCGA mesothelioma and PM tissue samples was calculated using xCell [69].

### Candidate targets of validated long noncoding RNAs

PM tissue expression data was used to identify potential target genes and pathways of PM-exo enriched LncRNAs. The expression of validated LncRNAs was correlated with the expression of all expressed protein-coding genes in PM tissue samples. For each LncRNA, the target genes were ordered based on decreasing correlation with its expression, and the ordered list of genes was used for enrichment analysis using fgsea R package (https://bioconductor.org/packages/release/bioc/html/fgsea.html) and hallmarks of cancer sets (https://www.gsea-msigdb.org/gsea/msigdb/). Only hallmarks with adjusted p-value < 0.01 were reported.

## Acknowledgements

We sincerely thank Dr. Emanuela Felley-Bosco for providing samples originating from SDM90 and SDM134 tissues.

## Author information

### Ethics declarations

All procedures involving pleural mesothelioma primary cultures were approved by the Ethics Committee of the canton Zurich on 09/02/2021 with the approval number BASEC-Nr. 2020–02566.

